# Corticosterone enhances formation of non-fear but not fear memory during infectious illness

**DOI:** 10.1101/2023.02.07.526836

**Authors:** Alice Hill, Colin Johnston, Isaac Agranoff, Swapnil Gavade, Joanna Spencer-Segal

**Author notes:** **Correspondence:** Joanna Spencer-Segal, MD, PhD.

## Abstract

Survivors of critical illness are at high risk of developing post-traumatic stress disorder (PTSD) but administration of glucocorticoids during the illness can lower that risk. The mechanism is not known but may involve glucocorticoid modulation of hippocampal- and amygdalar-dependent memory formation. In this study, we sought to determine whether glucocorticoids given during an acute illness influence the formation and persistence of fear and non-fear memories from the time of the illness. We performed cecal ligation and puncture in male and female mice to induce an acute infectious illness. During the illness, mice were introduced to a neutral object in their home cage and separately underwent contextual fear conditioning. We then tested the persistence of object and fear memories after recovery. Glucocorticoid treatment enhanced object discrimination but did not alter the expression of contextual fear memory. During context re-exposure, neural activity was elevated in the dentate gyrus irrespective of fear conditioning. Our results suggest that glucocorticoids given during illness enhance hippocampal-dependent non-fear memory processes. This indicates that PTSD outcomes in critically ill patients may be improved by enhancing non-fear memories from the time of their illness.

## 1 Introduction

More than one in five survivors of critical illness develop post-traumatic stress disorder (PTSD), exhibiting symptoms even up to eight years after discharge (Kapfhammer *et al*., 2004; Davydow *et al*., 2008; Bienvenu *et al*., 2018; Hatch *et al*., 2018). Glucocorticoids are often administered to critically ill patients, and several studies have found that this exposure to high levels of glucocorticoids during illness is associated with lower levels of PTSD after recovery (Schelling *et al*., 2001, 2004, 2006; Amos, Stein and Ipser, 2014; Sijbrandij *et al*., 2015). How glucocorticoids influence post-traumatic stress outcomes in critically ill patients is not known. Understanding the mechanisms underlying glucocorticoid modulation of PTSD risk will be crucial for informing critical illness treatment and improving long-term patient outcomes.

PTSD symptoms are characterized by pathological memories of the traumatic experience, leading to intrusions, avoidance behavior, and hypervigilance (Schelling *et al*., 2004; Hauer *et al*., 2009). Fragmentation of episodic memory, persistent fear memory, and fear generalization are specific memory processes that contribute to the development of PTSD. While the amygdala and hippocampus are both important for contextual fear, the hippocampus is central in controlling episodic memory and context recognition, ultimately enabling accurate recall of the experience and future discrimination between similar experiences or cues (Pennartz *et al*., 2011; Yassa and Stark, 2011).

The hippocampus and amygdala are both sensitive to glucocorticoids (Meijer, Buurstede and Schaaf, 2019), and glucocorticoids are known to modulate fear and non-fear memories. We sought to determine whether glucocorticoids given during an acute illness act on the hippocampus and amygdala to influence the formation and persistence of memories from the time of the illness.

Using a mouse model of systemic infection, cecal ligation and puncture, we tested the effects of corticosterone administration on long-term object memory and contextual fear memory by analyzing behavior and neural activity. We found that glucocorticoids given during illness specifically modulate non-fear memories in survivors, supporting an increased focus on the importance of non-fear memory for trauma outcomes.

## 2 Materials and Methods

### 2.1 Animals

Young adult 10-12-week-old C57BL6 male and female mice were obtained from the Jackson Laboratory (N = 80 total, half female). Animals were group housed on a 14:10 light/dark cycle with free access to food and water. All experimental protocols were approved by the University of Michigan Institutional Animal Care and Use Committee and conducted in accordance with the NIH Guide for the Care and Use of Laboratory Animals.

### 2.2 Induction of illness: cecal ligation and puncture

Animals were anesthetized with isoflurane and were injected with 60 uL of 0.25% bupivacaine at the incision site. Under aseptic conditions, a 5mm incision was made through the abdominal wall. The cecum was ligated 5mm from the end with a silk suture and then punctured through-and-through with a 19-gauge needle. The incision was closed with sutures. Surgery was immediately followed with subcutaneous injections of 1 mL saline and 0.1 mL imipenem-cilastatin. The day of surgery is referred to as Day 0. **Figure 1** shows the experimental timeline. As we were interested in the effects of glucocorticoid treatment on memory during illness, rather than the effects of illness itself, all animals underwent surgery.

**Figure 1.**
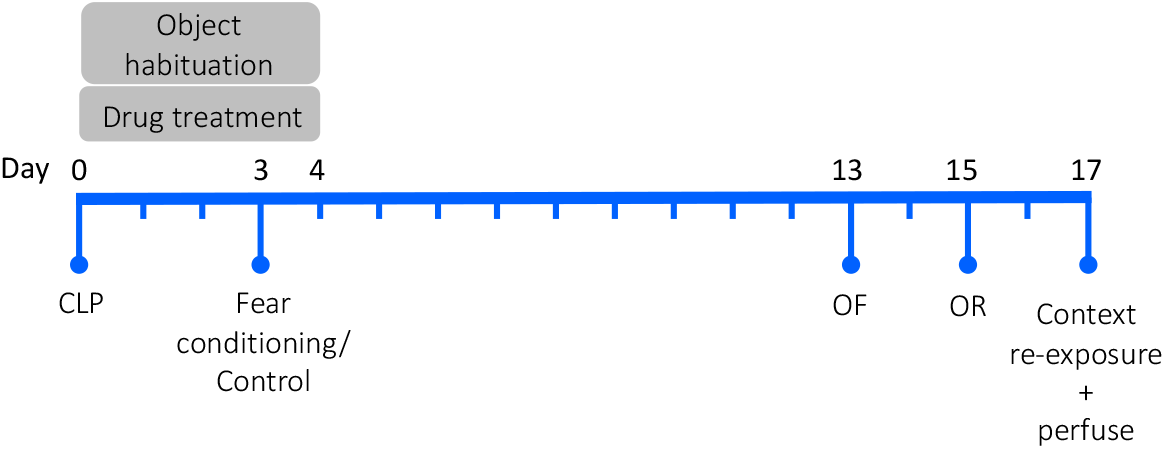
Experimental timeline. CLP = cecal ligation and puncture; OF = open field; OR = object recognition. N = 80 animals (20 animals/group, half female).

### 2.3 Drug treatment during critical illness

Animals received daily subcutaneous injections of 16 mg/kg corticosterone or vehicle on Days 0-4. The vehicle was 0.1 mL sesame oil.

### 2.4 Quantification of behavior after recovery from illness

Beginning on Day 13, after full physiologic recovery, animals underwent behavioral testing. Anxiety-like behavior was assessed by measuring exploration in a novel open field, episodic memory was assessed with a novel object recognition paradigm, and fear memory was assessed with a contextual fear-conditioning paradigm. All behavioral tests, except fear conditioning, were recorded and tracked using Ethovision software (version 15).

#### 2.4.1 Open field test

On Day 13, animals were placed in a novel brightly-lit open field (72 × 72 × 26 cm; 200 lux) and allowed to explore for 5 minutes.

#### 2.4.2 Novel object recognition

During the concurrent illness period and drug treatment period (Days 0-4), animals were habituated to one of two objects (counterbalanced across groups) in their home cage. On Day 15, after recovery, animals underwent a 5-minute object recognition discrimination trial under low-light conditions (30 lux) with two objects: the habituated object and a novel object. Exploration of each object was recorded, with the perimeter of the object defined as the object zone.

#### 2.4.3 Contextual fear-conditioning

During the concurrent illness period and drug treatment period (on Day 3), half the animals were subjected to a single footshock training session, consisting of 20 random one-second 0.45 mA shocks over 30 minutes. Olfactory and visual context was provided by 2% acetic acid and a blue-striped curtain. Control animals were placed in the same context but did not receive shocks. On Day 17, after recovery, animals were returned to the fear-conditioned context for 5 minutes and time spent freezing was recorded using FreezeFrame (ActiMetrics, Wilmette, IL).

### 2.5 Tissue collection

Two hours after testing in the fear-conditioned context on Day 17, all mice were injected with 0.1 mL Nembutal perfused with 0.1 M phosphate buffer (PB) followed by 4% paraformaldehyde (PFA). Brains were post-fixed with 4% PFA then 30% sucrose and stored at -80°C for future use.

### 2.6 Quantification of neural activity

Coronal brain sections (40 um) were cut serially on a sliding microtome (Leica Biosystems #SM2010R, Buffalo Grove, IL). Sections were rinsed in tris-buffered saline (TBS) and incubated overnight with 1:2000 rabbit primary alpha-mouse c-Fos antibody (SySy #226 003, Goettingen, Germany) in a buffer of bovine serum albumin (BSA), triton, and TBS. After several washes in TBC, sections were incubated with 1:300 biotinylated goat anti-rabbit IgG antibody (Vector Labs #BA-1000, Burlingame, CA) in a buffer of BSA and TBS for 30 minutes. This was followed by a 30-minute incubation in avidin-biotin-peroxidase complex (Vectastain Elite ABC-HRP Kit #PK-6100, Vector Labs, Burlingame, CA). Detection of antibody signal was revealed with 3,3’-diaminobenzidine (DAB) peroxidase substrate (DAB Substrate Kite, Peroxidase with Nickel, Vector Labs #SK-4100, Burlingame, CA).

DAB labeling was visualized using bright-field microscopy. Images were acquired on a Leica DMR-HC microscope (Leica Microsystems), with 5x objective using the same light conditions and acquisition parameters across all animals. In ImageJ, images were processed to remove background noise. The region of interest was selected with a standard-width ribbon tool. Area and automated cell count of the region were recorded. Relative c-Fos+ cell density was calculated by dividing the number of c-Fos+ cells by the area. This analysis was repeated for the basal amygdala and for dorsal and ventral sections of the CA1, CA3, and dentate gyrus. The endpoint for the histology was relative c-Fos+ cell density.

### 2.7 Data analysis

A three-way ANOVA revealed that shock history did not have a significant effect on open field and object recognition behavior, and so the effects of corticosterone and sex on open field behavior were compared using two-way ANOVA. For the object recognition test, several proportions were calculated: the proportion of total object exploration time spent with the familiar object and the proportion of total object exploration time spent with the preferred object. Animals with insufficient total object exploration (less than three seconds) were excluded from this following analysis (N = 9 animals). As shock history and sex had no effect on object recognition, the effect of corticosterone on novel object discrimination was compared to the chance value (50%) using an unpaired t-test in each of the corticosterone and vehicle groups. As sex also had no effect on freezing behavior, the effects of corticosterone treatment and shock history on freezing behavior were compared with a two-way ANOVA. Statistical analyses were performed using GraphPad Prism 9, with P < 0.05 considered significant. Graphs were generated in Prism and show individual data points plus the mean and standard error of the mean (SEM).

For c-Fos immunohistochemistry, a series of multiple linear regression models were generated to predict c-Fos+ cell density in the basal amygdala. Different combinations of the following predictors were included: sex, corticosterone treatment, and history of footshock. These models were compared against a null model and against each other using AICc. For each of the regions of interest in the hippocampus (CA1, CA3, dentate gyrus), a series of general multiple linear regression models were generated to predict c-Fos+ cell density, each including a random intercept to control for individual variation. Different combinations of the following predictors were included: sex, corticosterone treatment, history of footshock, and axis (ventral vs dorsal). The models were compared against a null model and against each other using AIC. These analyses were performed in R using the lme4 and bbmle packages. Graphs were generated in Prism and show individual data points plus the mean and standard error of the mean (SEM). Plots of the predictor effect sizes were generated using the sjPlot package in R.

## 3 Results

We used the gold standard mouse model of systemic infection, cecal ligation and puncture, to study the effects of glucocorticoid treatment during acute infectious illness on the long-term persistence of memories formed during illness (CLP; Buras, Holzmann and Sitkovsky, 2005). A total of 65 of the original 80 (81.25%) mice survived their infectious illness to undergo behavioral testing. Corticosterone treatment did not significantly improve survival; 35 of the 40 (87.5%) corticosterone-treated animals survived, while 30 of the 40 (75%) control animals survived (Fisher’s Exact Test; Relative risk: 0.857, P = 0.252).

### 3.1 Corticosterone had no effect on anxiety-like behavior

We previously showed that mice begin physiologic recovery from CLP within 5 days, with complete recovery of body weight and locomotion within 14 days (Spencer-Segal *et al*., 2020). Here, we assessed affective behavior in 14-day CLP survivors with an open field test in which mice explored a mildly aversive brightly-lit novel arena. In our prior study, we found that 2-week CLP survivors showed negative affective behavior as compared to a sham operation, and here we asked whether glucocorticoid treatment during illness influences this behavior in CLP survivors. There was a sex difference in open field behavior: females traveled further (F(1,61) = 25.41, P < 0.001), spent longer in the center (F(1,61) = 7.547, P = 0.008), and entered the center more frequently than males (F(1,61) = 26.56, P < 0.001). We found that corticosterone treatment did not impact exploration in the open field in either sex; corticosterone-treated mice did not differ from vehicle-treated mice in total distance traveled (**Fig. 2A**; F(1,61) = 0.004, P = 0.951), duration in the brightly-lit center (**Fig. 2B**; F(1,61) = 0.496, P = 0.484), or frequency in the center (**Fig. 2C**; F(1,61) = 0.494, P = 0.485). Thus, corticosterone treatment during illness did not affect behavior in the open field after recovery from illness.

**Figure 2.**
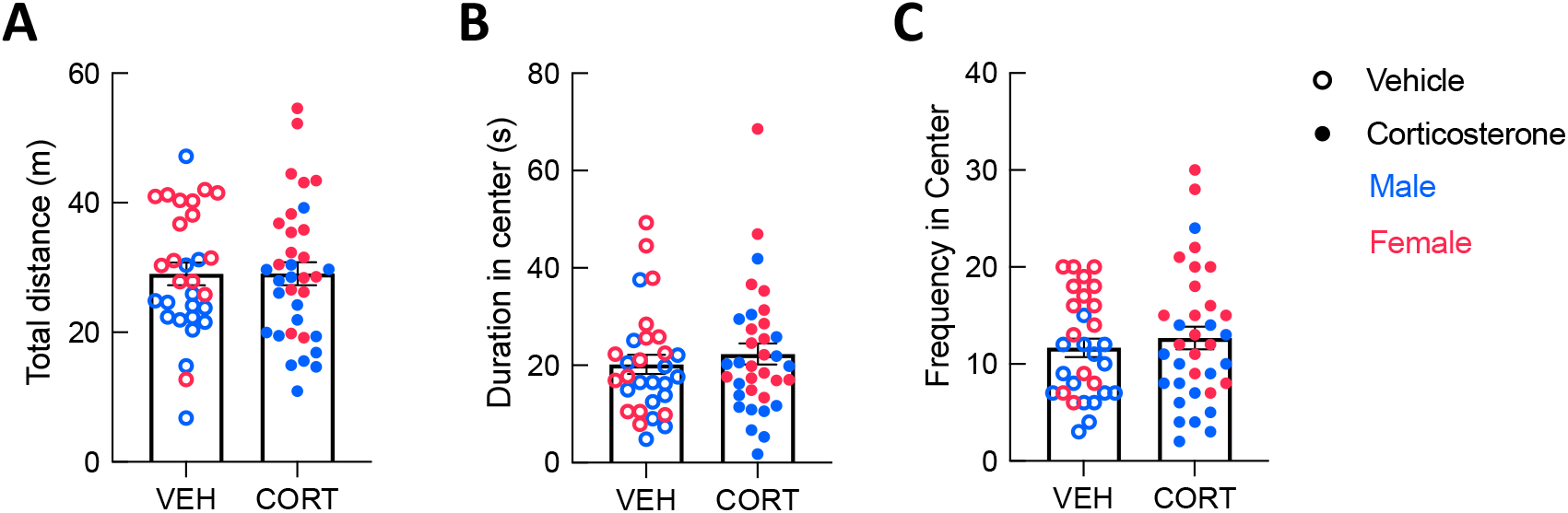
Corticosterone administration did not affect locomotion or anxiety-like behavior in the open field. There was no difference in (A) total distance traveled, (B) total time spent in the brightly-lit center, and (C) the number of entries into the center between corticosterone-treated and control animals. In all three measures, females were more active than males. (VEH: N = 30; CORT: N = 35).

### 3.2 Corticosterone facilitated memory of an object from illness

We tested the effect of corticosterone on non-fear memory with a novel object recognition paradigm in which animals were introduced to an object in their home cage during their illness. After recovery, memory of the object from illness was assessed using a discrimination trial in which the animal could freely explore this familiar object or a novel object inside a familiar arena.

In the discrimination trial, corticosterone-treated mice showed no difference in locomotion in the arena (**Fig. 3A**; F(1,61)=1.557, P=0.217). In contrast, corticosterone did influence overall object exploration: corticosterone-treated animals spent more total time exploring the objects than vehicle-treated animals (**Fig. 3B;** t(63) = 2.011, P = 0.049). Corticosterone-treated mice also tended to spend more time with the familiar object than did those treated with vehicle (t(63)=1.982, P=0.052), but there was no effect of corticosterone treatment on time spent exploring the novel object (t(63) = 1.138, P = 0.260). This suggests that animals treated with corticosterone selectively explored the familiar object over the novel object. Indeed, corticosterone-treated animals exhibited a significant preference for the familiar object (t(29) = 2.599, P = 0.015), while vehicle-treated animals had no preference (**Fig. 3C**; t(25) = 0.897, P = 0.378). The discrimination between the novel and familiar object in corticosterone-treated mice suggests that corticosterone-treated mice recalled a long-term memory of the object last seen 11 days prior, while vehicle-treated animals did not.

**Figure 3.**
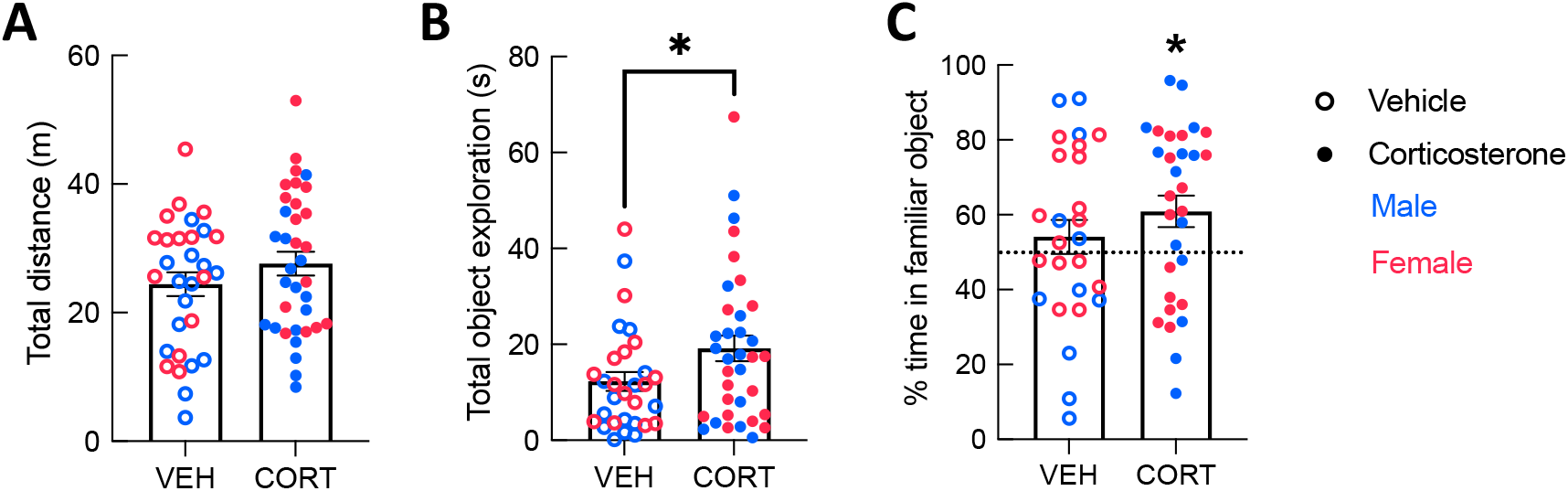
Corticosterone-treated mice showed improved object recognition. Corticosterone-treated animals exhibited (A) similar levels of overall activity as control animals but (B) spent more time exploring the objects. Corticosterone-treated animals demonstrated a preference for the familiar object over the novel object (VEH: N = 30; CORT: N = 35). *P < 0.05.

### 3.3 Corticosterone did not affect fear memory from illness

To assess the effects of corticosterone on fear memory persistence after illness we used a contextual-fear conditioning paradigm in which animals underwent a single training trial during the illness period. Two weeks later, after recovery, animals were re-exposed to the conditioned context in a single testing session and freezing behavior was recorded to measure fear memory.

Corticosterone treatment did not affect freezing during training (**Fig. 4A**; F(1,53) = 0.297, P = 0.588), which was higher in the shock than in the no-shock group (F(1,53) = 7.633, P = 0.008). During retention testing, fear-conditioned mice showed increased freezing in the conditioned context (**Fig. 4B**; F(1,61) = 128.9, P < 0.001), demonstrating that all animals formed a lasting fear memory, with no effect of corticosterone (F(1,61) = 0.350, P = 0.557). Corticosterone therefore had no effect on fear memory in this assay, in contrast to its facilitation of object memory.

**Figure 4.**
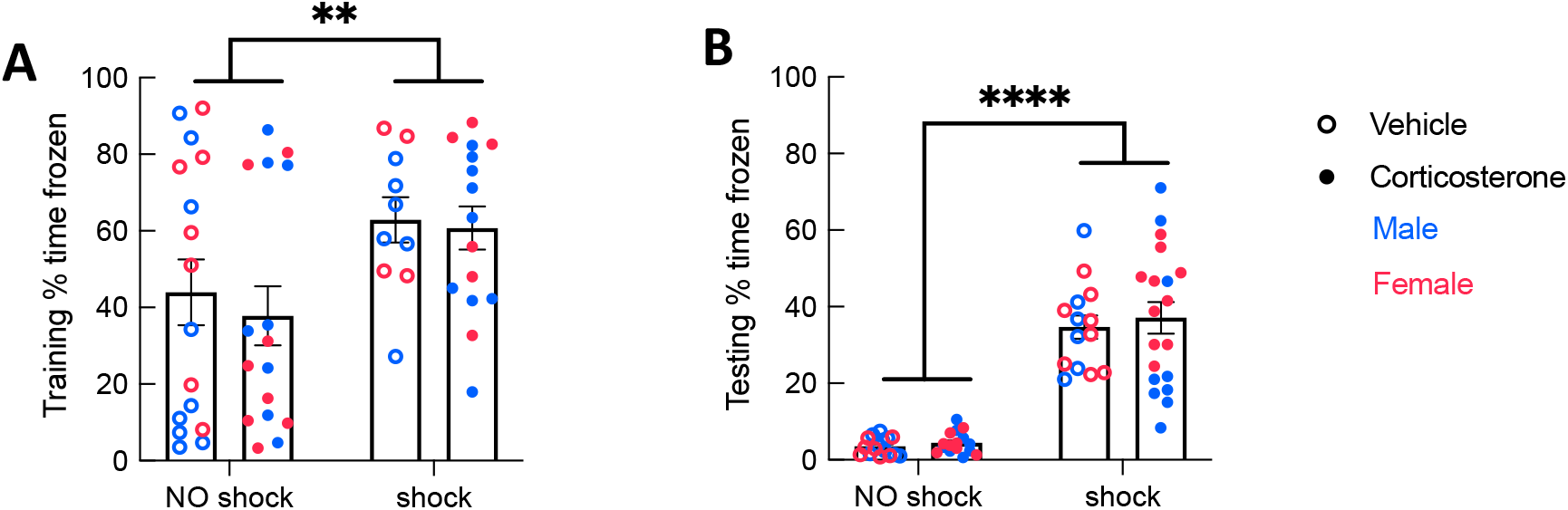
History of footshock, but not corticosterone administration, affected contextual fear memory. Animals that had experienced shocks from more than control animals in both (A) training and (B) testing trials, while corticosterone-treated and vehicle-treated animals froze at equal rates (Training: No shock N = 32, Shock N = 25; Testing: No shock N = 32, Shock N = 33). *P < 0.05; * *P < 0.01; * * * * P < 0.0001.

### 3.4 Corticosterone treatment enhanced neural activity associated with a familiar context

After the context re-exposure during the testing trial, mice were euthanized for quantification of neural activity in the hippocampus and basal amygdala using c-Fos immunohistochemistry as a proxy for neuronal activation in those brain regions (**Figure 5**). Significant predictors of c-Fos+ cell density were identified via linear regression and model comparison. In the CA1 and CA3, there was no significant contribution of corticosterone treatment and shock (**Fig. 5A and B**). The top model for both regions included only axis as a predictor, indicating that neural activity was higher in the ventral area (**Table 1**). The models that included corticosterone treatment and shock as predictors did not fit the data as well, and the effect sizes for these predictors were not significant. In the dentate gyrus, animals treated with corticosterone had greater c-Fos+ cell density compared to animals treated with vehicle (**Fig. 5C**; P = 0.002), suggesting greater activation of the dentate gyrus during context re-exposure in the corticosterone-treated mice. The top model for the dentate gyrus included corticosterone as a predictor and the effect size was significant (**Table 1**). In all hippocampal regions, there was no interaction between the effect of corticosterone treatment and the effect of history of footshock on neural activity; the models that included an interaction between corticosterone treatment and shock fit the data poorly and the effect size of the interaction term was not significant (**Table 1**).

**Table 1.**
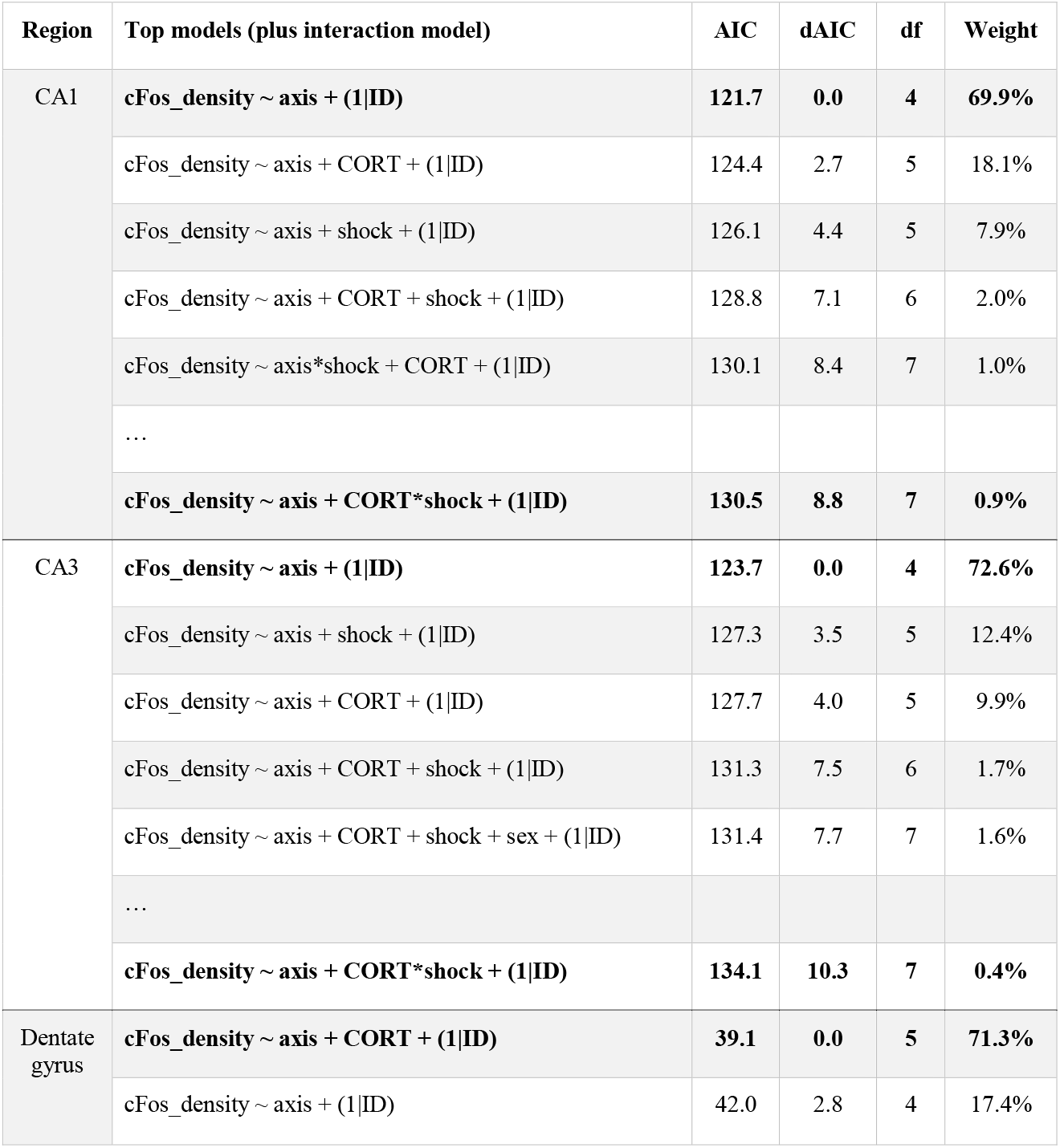
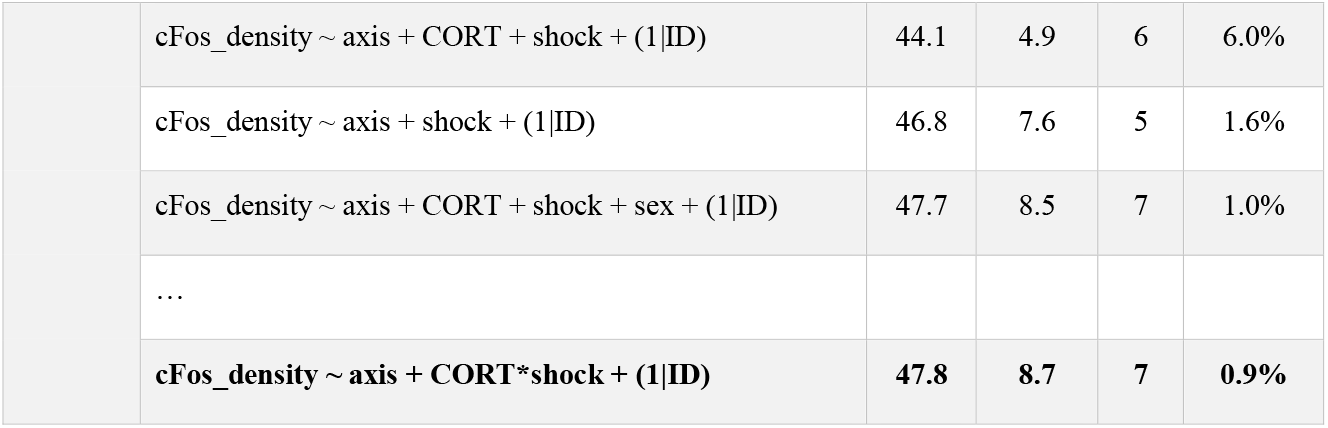
Model selection for the effect of axis (ventral), history of footshock (shock), corticosterone treatment (CORT), and sex on c-Fos+ cell density in the hippocampus. General linear models using different combinations of predictors and controlling for individual variation (ID) were fit for each of the three hippocampal brain regions separately. For each region, the top 5 models are displayed, in addition to the interaction model (which reliably fit the data poorly). Lower AIC values indicate better fitting models. Only in the dentate gyrus did the top model include corticosterone treatment as a significant predictor of neural activity.

**Figure 5.**
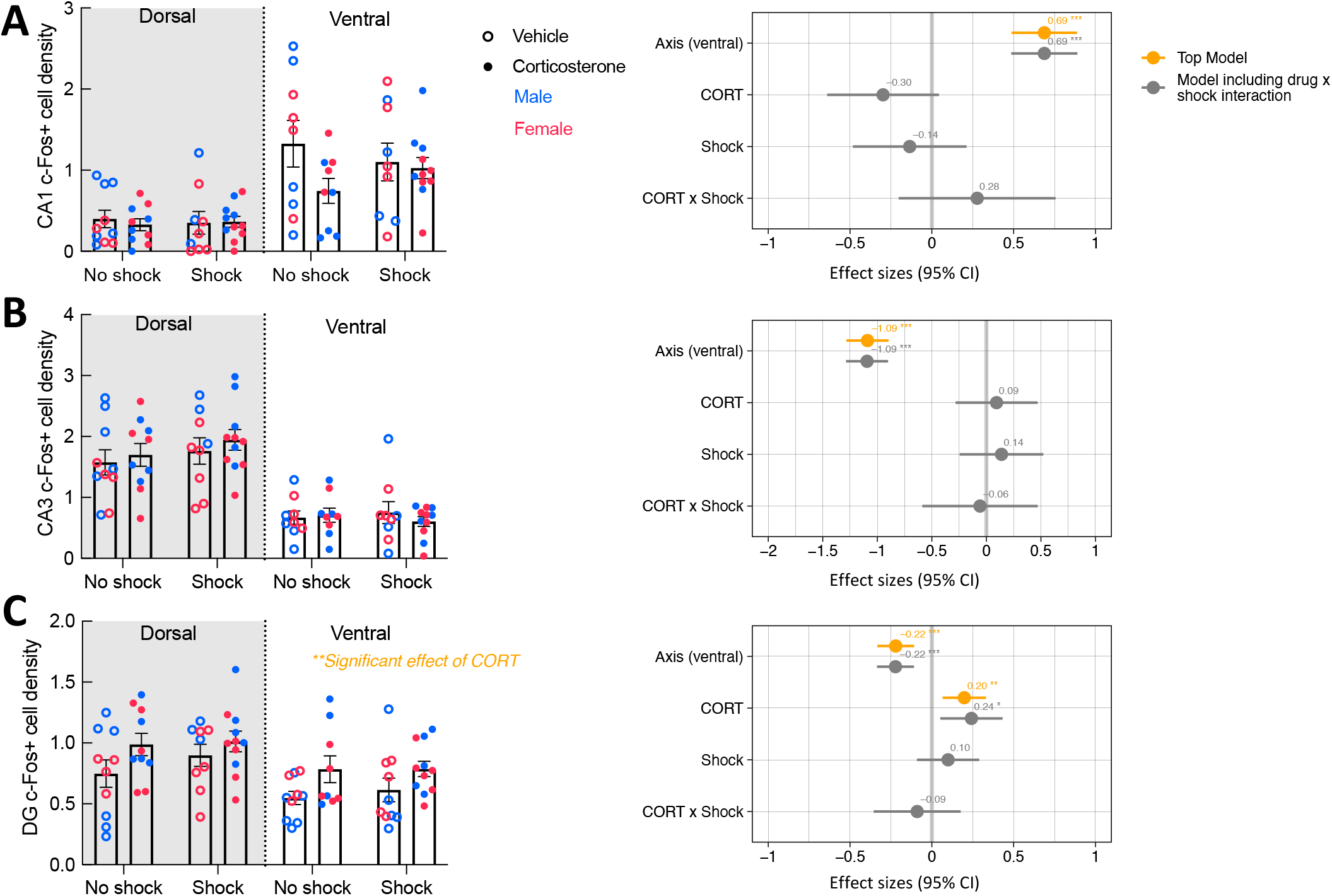
Corticosterone treatment increased neural activity in the dentate gyrus during context re-exposure. The density of c-Fos+ cells in the (A) CA1 and (B) CA3 were not affected by history of footshock or corticosterone administration. (C) In the dentate gyrus (DG), c-Fos+ cell density was greater in corticosterone-treated animals than control animals. The forest plot on the right of each panel shows the effect sizes of the predictors with a 95% confidence interval. The orange points represent the effect sizes of the predictors in the top model, while the grey represents the predictor effect sizes of the model that includes the CORT x Shock interaction term. Only in the dentate gyrus does corticosterone treatment appear as a predictor in the top model, with a significant positive effect on neural activity. * *P < 0.01; * * * P < 0.0001.

While fear conditioning (shock vs. no shock) did not affect c-Fos+ cell density in the hippocampus, it did so in the basal amygdala, where fear conditioned mice did show increased c-Fos+ cell density (**Fig. 6A**; P = 0.038). The top model included only history of footshock as a predictor, and the effect size was significant (**Table 2**). There was no effect of corticosterone treatment on c-Fos+ in the amygdala and again, there was no significant interaction between the effect of corticosterone treatment and history of footshock (**Fig. 6B**). This is indicated by the poorly fitting interaction model, in which the effect size of the interaction term was not significant (**Table 2**). In conclusion, c-Fos+ immunohistochemistry demonstrated increased neuronal activation in the dentate gyrus during context re-exposure in corticosterone-treated mice, independent of fear conditioning.

**Table 2.** Model selection for the effect of history of footshock (shock), corticosterone treatment (CORT), and sex on c-Fos+ cell density in the basal amygdala. Linear models using different combinations of predictors were compared using AICc, due to small sample size. The poorly fitting interaction model is also included for reference. Lower AICc values indicate better fitting models. The top model includes history of shock as a significant positive predictor of neural activity in the basal amygdala.

| Top 5 models (plus interaction model) | AICc | dAICc | df | Weight |
| --- | --- | --- | --- | --- |
| **cFos_density ~ shock** | **-4.6** | **0.0** | **3** | **46%** |
| cFos_density ~ 1 (null model) | -2.3 | 2.2 | 2 | 15.1% |
| cFos_density ~ CORT + shock | -2.0 | 2.5 | 4 | 12.9% |
| cFos_density ~ CORT + shock + sex | -1.7 | 2.8 | 5 | 11.1% |
| cFos_density ~ CORT | 0.1 | 4.6 | 3 | 4.6% |
| ... |  |  |  |  |
| **cFos_density ~ CORT\*shock** | **0.7** | **5.2** | **5** | **3.4%** |

**Figure 6.**
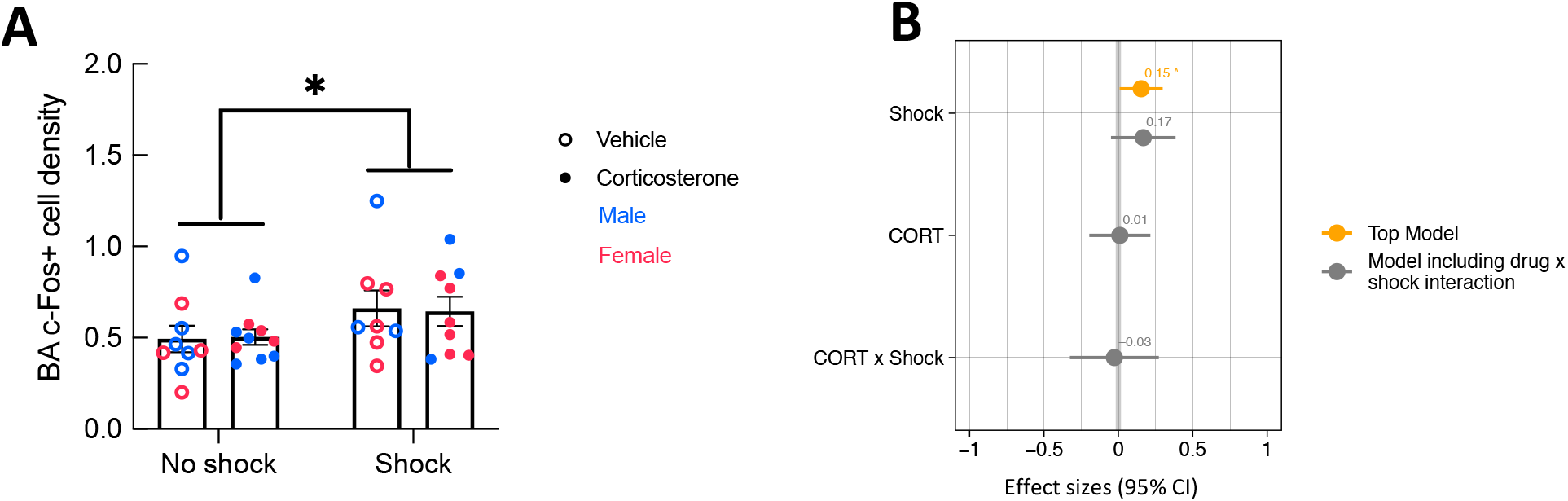
History of footshock, but not corticosterone treatment, increased neural activity in the basal amygdala during context re-exposure. (A) The density of c-Fos+ cells was greater in animals previously exposed to footshocks than in control animals. (B) the top linear regression model included only history of footshock as a predictor, with a significant positive effect size. The model including an interaction term between corticosterone treatment and shock was a worse fit and included no meaningful effect of the interaction or of each independent predictor. *P < 0.05.

## 4 Discussion

We found that glucocorticoid treatment during illness impacted the persistence of non-fear memory but not fear memory after recovery. Administration of corticosterone during CLP-induced illness facilitated long-term memory of an object from the illness period. In contrast, neither affective behavior nor the expression of contextual fear memory was affected by corticosterone treatment. Corticosterone treatment was also associated with elevated neural activity in the dentate gyrus during context re-exposure independent from fear conditioning. This increase in dentate gyrus neural activation during context re-exposure suggests that corticosterone promoted memory of the context, rather than fear memory per se. Taken together, our data suggest that corticosterone given during illness primarily impacts hippocampal-dependent non-fear memory processes. In the context of the repeated finding, in humans, that glucocorticoid treatment during illness decreases the risk of PTSD in survivors, our data suggest the intriguing possibility that glucocorticoids treatment prevent PTSD by improving episodic memories from the illness.

As mice are assumed to prefer novelty, we were surprised to see that the animals spent more time with the familiar than the novel object during the object discrimination test. While unusual, the apparent behavioral discrimination still suggests intact memory of the familiar object. The preference for the familiar object could be due to a positive emotional valence attached to the object during illness. It could also be due to the fact that the discrimination trial was run in a familiar arena while mice had previously seen the object in their home cage, leading to recognition of an object-context mismatch during the discrimination trial.

Contextual fear memory depends on both the hippocampus and the amygdala. The amygdala, particularly the basal amygdala, is associated with the emotional aspect of the memory, while the hippocampus pairs the fear memory to the context (Phillips and LeDoux, 1992; Kim and Cho, 2020). In animal studies, targeted administration of glucocorticoids to the basal amygdala enhances its activity and fear memory consolidation and acquisition (Roozendaal and McGaugh, 1997; Donley, Schulkin and Rosen, 2005; Roozendaal, McEwen and Chattarji, 2009; Finsterwald and Alberini, 2014), whereas general administration of glucocorticoids, presumably available to both the hippocampus and the amygdala, results in impaired fear memory consolidation and retrieval (de Quervain, Roozendaal and McGaugh, 1998; Cohen *et al*., 2008).

We found that corticosterone did not affect fear memory after conditioning during CLP-induced illness. All animals exposed to shocks showed good retention, which was not altered by corticosterone treatment. It is possible that in our study we observed a “ceiling effect” considering the high efficacy of our fear conditioning regiment in both corticosterone-treated and untreated groups. But the findings are consistent with several clinical studies that find that the preventative effect of glucocorticoids against PTSD is not associated with reduced numbers or types of fear memory (Schelling *et al*., 2001, 2004; Weis *et al*., 2006). It is also possible that the effects of glucocorticoids on fear memory were not captured with our paradigm. Other studies have found that PTSD is marked by abnormal extinction recall and fear renewal (Garfinkel *et al*., 2014); perhaps if we had tested the animals for extinction, extinction recall, and renewal of the fear memory, we would have seen an effect of glucocorticoids on behavior and amygdala activity.

C-Fos cell density analysis revealed greater neural activity in the basal amygdala in the fear-conditioned group, supporting the role of amygdala in fear memory recall and consistent with the lack of effect of corticosterone on freezing behavior during the testing trial. In contrast, animals treated with corticosterone exhibited greater neural activity in the dentate gyrus during context re-exposure, irrespective of fear conditioning. As granule cells in the dentate gyrus support context discrimination and memory recall (Hainmueller and Bartos, 2020), this potentially indicates improved recognition of the testing context in corticosterone-treated mice, consistent with the above improvement in episodic memory from the illness period. Consistent with a possible role for the dentate gyrus in the lasting effects of glucocorticoid treatment, a previous study showed that hydrocortisone given to stress-exposed animals caused lasting increases in dendritic growth and spine density in dentate granule cells (Zohar *et al*., 2011). The dentate gyrus has previously been implicated in the stimulus overgeneralization and subsequent inappropriate emotional responses seen in disorders such as PTSD (reviewed in Kheirbek et al., 2012). Our findings suggest that glucocorticoids may act at the dentate to improve recall of non-fear memories from illness and decrease PTSD risk.

Other brain regions previously implicated in stress-associated memories include the medial prefrontal cortex (mPFC). While the hippocampus encodes memories of the context, the PFC is responsible for associating these memories with emotional responses (Euston, Gruber and McNaughton, 2012). The two regions demonstrate strong connectivity and are functionally linked (Cenquizca and Swanson, 2007; Herweg *et al*., 2016). Like the hippocampus, there is high glucocorticoid sensitivity in the mPFC and, in fact, expression of contextual fear conditioning depends upon activation of glucocorticoid receptors in the mPFC (Reis *et al*., 2016). In addition, diminished mPFC signaling has been repeatedly associated with PTSD (Etkin and Wager, 2007). In the future, it would be interesting to investigate the effect of glucocorticoid treatment on mPFC activity during recall of fear and non-fear memories from illness.

Taken together, our findings suggest that corticosterone treatment during illness improves non-fear memory from the illness period without affecting fear memory. Accurate recall of the ICU experience (rather than delusional or emotional memories) has been proposed to protect against PTSD in survivors (Jones *et al*., 2001); this is the presumed mechanism for clinical interventions such as ICU diaries (Jones *et al*., 2010; Garrouste-Orgeas *et al*., 2012, 2019; Wang *et al*., 2020). Our findings suggest that glucocorticoids may act via this mechanism during illness and suggest that continued focus on enhancing episodic memories from the illness period may be valuable PTSD prevention strategies in critically ill patients and survivors.

## 7 Author Contributions

JSS and AH applied for grants, designed the experiment, and wrote the manuscript. AH created the figures. AH, CJ, and IA performed the experiments. SG contributed to immunohistochemistry data analysis. All authors read and agreed to the published version of the manuscript.

## 9 Contributions to the field

Glucocorticoids given during critical illness are known to decrease the risk of post-traumatic stress disorder (PTSD) in survivors, but how they do this is unknown. Since memories of the traumatic experience contribute to PTSD, we studied the effect of giving glucocorticoid treatment during acute illness in mice on long-term memories formed during the illness. We found that glucocorticoids did not affect fear memory, but instead improved non-fear memory and enhanced the neural representation of a contextual memory. The results suggest that glucocorticoids given during critical illness specifically modulate non-fear memories. Strengthening these memories from the time of illness may be a viable PTSD prevention strategy in critical illness survivors.

